# Biogeographical patterns and diversity in the diet of the culpeo (*Lycalopex culpaeus*) in South America

**DOI:** 10.1101/2021.09.09.459655

**Authors:** Jorge Lozano, Marta Guntiñas, Rodrigo Cisneros, Esther Llorente, Aurelio F. Malo

**Author notes:** Correspondence Author, Jorge Lozano, Departamento de Biodiversidad, Ecología y Evolución, Universidad Complutense de Madrid, C/ José Antonio Novais 12, 28040, Madrid, Spain. These authors contributed equally to this manuscript (Jorge Lozano, Marta Guntiñas).

## Abstract

Here we describe the dietary patterns of the culpeo (or Andean fox) at a biogeographical scale. We also analyze the influence of exotic lagomorphs on its diet and explore differences between culpeo subspecies. We selected 17 mutually comparable diet studies, which include 19 independent diet assessments. Then, we extracted and standardized the values of the different diet components from these studies and calculated the relative frequency of occurrence (RF) of the ten main trophic groups that we found. Further, we calculated the Shannon-Wienner H’ trophic diversity index.

The results showed that small mammals (41%), lagomorphs (21%), invertebrates (12.4%) and large herbivores (7.3%) were the most consumed groups. A factorial analysis of all trophic groups rendered four orthogonal factors that were used as response variables in relation to a set of environmental predictors. Altitude correlated with most factors (i.e. trophic groups). Exotic lagomorphs were consumed in lowlands, in higher latitudes and in regions showing high values of the human footprint index, replacing in these areas native fauna as the main prey. There were no differences in diet between the two main culpeo subspecies analysed, *L*.*c. culpaeus* and *L*.*c. andinus*.

Finally, the best explanatory models (GLM) of trophic diversity selected, using the Akaike’s information criterion (AIC), showed that the most diverse diets were those composed of large herbivores, edentates, carnivorous species, birds and herptiles (i.e. amphibians and reptils), in areas of high rainfall located in protected areas. Neither latitude nor altitude seemed to have an effect on the trophic diversity of the culpeos, as they were not retained by the final models.

## Introduction

The functioning and diverse characteristics of ecosystems are determined and conditioned by a multitude of ecological processes and parameters, such as evolutionary history and the dynamics of species interaction, such as parasitic, competitive or predator-prey relationships, the latter being the most important in the configuration of food webs (Ritchie et al., 2012; Soe et al., 2017). In fact, large carnivores and mesopredator predation tend to play a prominent, and often key, role in the functioning of ecosystems (Newsome & Ripple, 2015), affecting prey population cycles (Kausrud et al., 2008) and producing multiple cascading effects (Coulson & Malo 2008; Ripple et al., 2014).

In turn, the ecology of a predatory species is influenced by the conditions of the environment in which it lives. Thus, when the different populations that make up a widely distributed species are considered, there is a wide range of environmental conditions that vary across its range. Faced with different ecological conditions each species presents a certain degree of flexibility in their behavior, which depending on its range and geneflow between subpopulations, can define the species as generalist or specialist. In particular, and in relation to the extent of the trophic niche and prey selection criteria, predators can be considered as generalists or opportunists, strict specialists or as facultative specialists (Glasser, 1984; Jaksic, 2007).

In the case of predators that have a wide distribution range, strong spatial variation in diet patterns is observed when studied at a biogeographic scale, as shown for numerous species such as the brown bear (*Ursus arctos*) (Vulla et al., 2009), the Eurasian otter (*Lutra lutra*) (Clavero et al., 2003), the European wildcat (*Felis silvestris*) (Lozano et al., 2006), the European badger (*Meles meles*) (Hounsome & Delahay, 2005), or the red fox (*Vulpes vulpes*) (Soe et al., 2017). For example, it is usual to find clear correlations in diet variation with latitude and climate, also related to the trophic plasticity detected with generalist or facultative specialist strategies (Virgós et al.,1999; Cox & Moore 2005; Lozano et al., 2006). Indeed, diet can have important implications at several levels. On the one hand, there are a multitude of factors associated with the trophic behavior of species, which can have repercussions on other aspects of their ecology. For example, the abundance and distribution of native prey not only affect dietary patterns but also space use, morphological characteristics and mating cycles (Dayan & Simberloff, 1996). On the other hand, the incorporation of exotic or non-native species into the diet and the consumption of anthropogenic food waste can also directly affect the abundance, distribution and behavior of carnivore species (Bino et al., 2010). In addition, processes of competition among predators and their differential vulnerability can modify both resource use patterns, as seen in medium-sized carnivores in North America (Lesmeister et al., 2015), and physical characteristics of the predator, such as body size, through processes of niche segregation (Jimenez et al., 1996).

To the factors mentioned above we must add the growing uncertainty on the ability of predators and prey to respond to the effects of global phenomena such as climate change (Bailey & van de Pol, 2016; Keith & Bull, 2016), and other anthropogenic factors (Sandom et al., 2014), which can promote important changes in trophic interactions at a global scale (Grosbois et al., 2008; Merilä, 2012) triggering profound changes in the structure of ecosystems (Newsome et al, 2015). For this reason, a better understanding of the trophic ecology of species and their interaction with environmental factors at various scales is necessary to develop species conservation strategies that mitigate the effects of global problems on ecosystems.

The way to evaluate the trophic strategy and plasticity of a species is to approach its study at large spatial scales, through the comparison of habitat- and region-specific diet within its range, based on the review and meta-analysis of local data. For many species this biogeographical dietary approach is not yet possible due to the few studies available at a local scale. Fortunately, this is not the case of the culpeo (*Lycalopex culpaeus*), also called Andean fox, a canid that is distributed in various habitats from southern Colombia, through Ecuador, Peru, Bolivia and Chile, to Patagonia and Tierra de Fuego in Argentina (Lucherini, 2016; Guntiñas et al., 2021), for which a number of diet studies have been published. Although on a global scale the International Union for Conservation of Nature (IUCN) classified the culpeo for its entire range as ‘Least Concern’ (Lucherini, 2016), this good conservation status is not mirrored at regional scales. For instance, in Colombia and Ecuador the culpeo is listed as a threatened species (Tirira, 2011; MADT, 2014), emphasizing the need to deepen the knowledge of its ecology.

Local studies on the culpeo diet interpret the trophic strategies of the species differently (see Guntiñas et al., 2021), considering it a strict carnivore (Iriarte et al., 1989; Redford & Eisenberg, 1992; Jiménez & Novaro, 2004), a practically insectivorous predator (Guzmán-Sandoval, 2007) with tendencies to frugivory (Ebensperger et al., 1991; Cornejo-Farfán & Jiménez-Milón, 2001), or a predator with high level of trophic plasticity that is capable of using a wide combination of resources (Jaksic et al., 1993; Johnson & Franklin 1994b; Castro et al., 1994). Recent studies suggest that culpeo behaves more like a facultative specialist (Guntiñas et al., 2017). However, to date no studies have been carried out that consider the data as a whole at a biogeographical scale so that the global trophic patterns can be detected, as well as their ecological correlates, which would allow a panoramic view of the culpeo trophic ecology to be obtained.

The present work reviews culpeo diet studies and data published throughout the species range, with the following aims: 1) to describe the general patterns of the culpeo diet at a biogeographical scale through a meta-analysis and the environmental factors that determine them; 2) to assess the degree of the consumption of exotic lagomorphs by culpeos and, in particular, the influence of this consumption on the use of native fauna as a trophic resource (Crespo & de Carlo, 1963; Jaksic, 1998; Novaro et al., 2000a; Rubio et al., 2013); 3) to evaluate the dietary differences between the two main culpeo subspecies described by Guzmán et al. (2009); and 4) to obtain an explanatory model of the trophic diversity of culpeo in South America.

## Material and Methods

We carried out a complete compilation of the published articles and other reports on the diet of the species. For information published prior to 1988 we used the review by Medel & Jacsik (1988). For works published after 1988, we carried out a systematic bibliographic search using the ‘Web of Knowledge’, ‘Google Scholar’ and ‘Scopus’ servers, including terms such as diet, culpeo, Andean fox, *Lycalopex culpaeus*, páramo wolf, and also the specific names that have been used to describe the culpeo previously (i.e. *Dusicyon culpaeus* and *Pseudalopex culpaeus*) (see for more details Guntiñas et al., 2021).

Once we selected all the existing studies, a subset was chosen to be used for statistical analyses. These studies met a number of requirements to ensure the representativeness of the data and the statistical power (Guntiñas et al., 2021). Firstly, they had to provide tables of data on the culpeo diet in the form of either frequency of occurrence, relative frequency of occurrence or number of prey items, as well as the sample sizes for the scats or stomachs analyzed. The data had to describe all the resources consumed by the species, so that studies with data focusing only on a particular prey group were excluded. Secondly, the papers had to have a minimum sample size of 30 (Lozano et al., 2006; Soe et al., 2017) per habitat type considered, treating these habitat units as independent samples. And finally, data had to be representative of the annual cycle in each habitat unit (Lozano et al., 2006).

With all, 17 (16 articles and one thesis) studies were finally selected meeting the aforementioned criteria, with a total of 19 independent samples (see Figure 1). Altogether the dataset included 4115 scats and/or stomachs with a total of 6220 trophic elements. Due to the variability in data presentation between studies, we standardized the data of the selected studies by calculating the relative frequency of occurrence (RF) for each independent sample. For the analyses, we did not consider whether identified trophic items came from stomachs or scats (Lozano et al., 2006).

**Figure 1.**
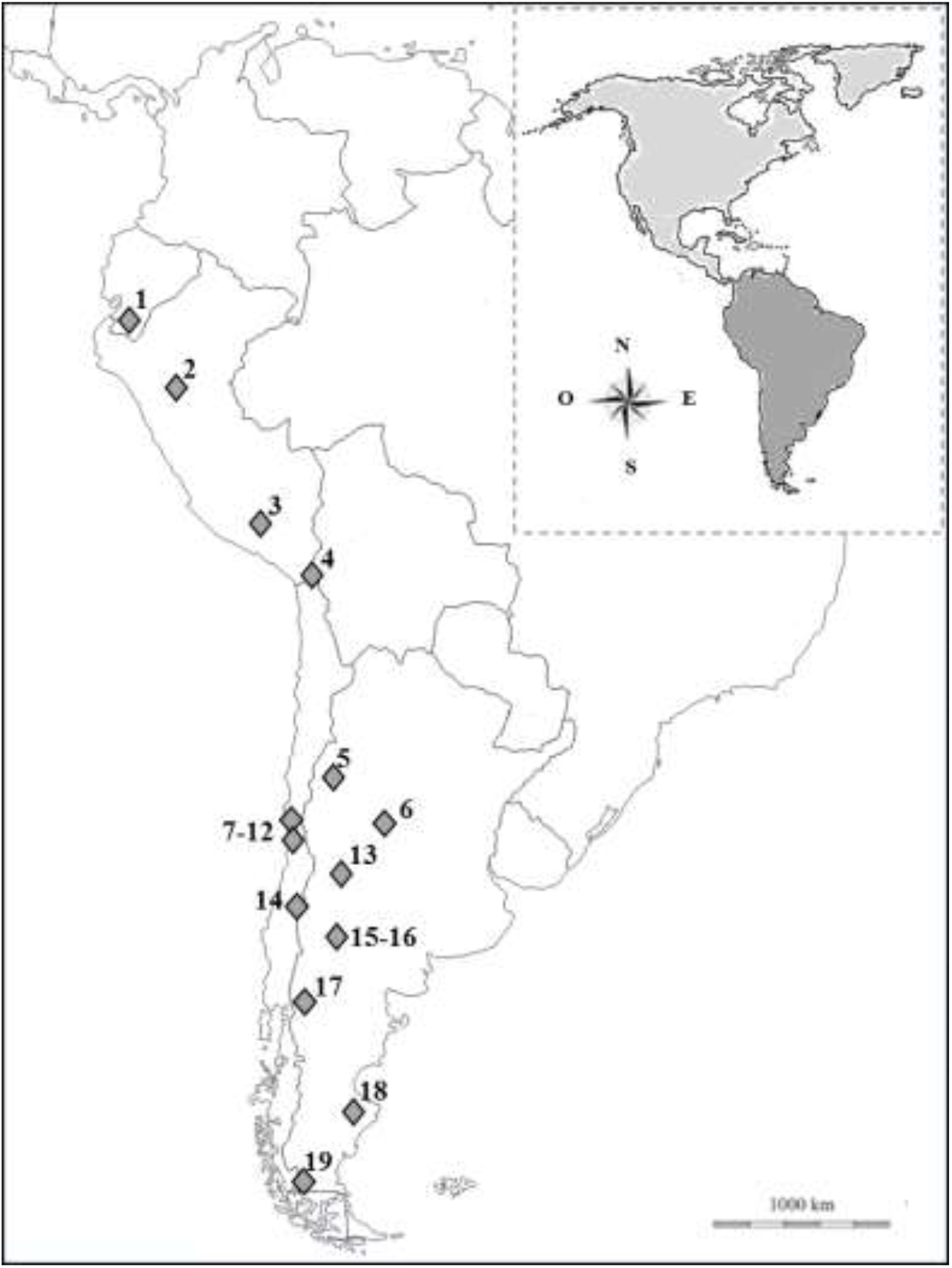
Location of the different study areas: 1. Guntiñas *et al*., 2017 2. Romo *et al*., 1995 3. Cornejo & Jiménez, 2001 4. Marquet *et al*., 1993 5. Walker *et al*., 2007 6. Pía *et al*., 2003 7. Ebensperger *et al*., 1991 8-9. Iriarte *et al*., 1989 10. Jaksic *et al*., 1980 11-12. Rubio *et al*., 2013 13. Berg, 2009 14. Achilles, 2007 15. Palacios *et al*., 2012 16. Novaro *et al*., 2000 17. Monteverde *et al*., 2011 18. Zapata *et al*., 2005 19. Johnson y Franklin, 1994

### Selection of variables

We categorized the different elements that composed the culpeo diet into ten main trophic groups (Guntiñas et al., 2021): ‘small mammals’ (i.e. mammals weighing less than 300 g, mainly rodents, insectivores and some marsupial species), ‘big rodents’, ‘lagomorphs’, ‘carnivorous’ (i.e. carnivores and carnivorous marsupials), ‘edentates’ (i.e. armadillos); ‘large herbivores’ (i.e. deer, camelids and livestock), ‘birds’, ‘eggs’, ‘herptiles’ (i.e. amphibians and reptiles), and ‘invertebrates’. The relative frequency of occurrence (RF) for each of these trophic groups was recalculated by dividing the number of items in each group by the total number of items. The presence of plant matter in the diet of each study was also recorded.

We also calculated the trophic diversity of the culpeo in each independent sample as the value of the Shannon-Wienner H’ diversity index (Weaver & Shannon, 1949). In order to calculate and compare the value of the index between the different studies, the trophic groups were standardized to the same taxonomic level as the groups in the less detailed publications, leaving the following 19 groups: rodents, soriids, procyonids, edentates, mustelids, marsupials, lagomorphs, cattle (sheep), cervids, camelids, felids, carrion, birds, reptiles, fish, amphibians, invertebrates, eggs and rubbish.

We recorded the main habitat type in which the work was carried out using the geographical location of each of the studies where the samples have been taken. Further, we determined whether the site was part of a protected area or not, the average annual rainfall, the altitude and the latitude. We also measured the degree of interference of human activity in the territory by calculating the human footprint index for each independent sample. To do it, we used two of the human footprint index layers which were downloaded (for the years 1993 and 2009) from the Center for International Earth Science Information Network (CIESIN). Based on the sampling year for each study, we attributed the value of the index layer corresponding to the nearest year to each area. On the other hand, we tested whether introduced (i.e. exotic) lagomorph species existed in the different study areas. For this purpose, the layers of presence of the species of interest available on the IUCN server were downloaded and the overlap with the sampling areas of the selected studies was checked. To observe possible differences in diet according to the subspecies, each study was attributed the subspecies *L*.*c. andinus, L*.*c. culpaeus* or *L*.*c. reissii* based on their geographical distribution (Guzmán et al., 2009; Guntiñas et al., 2021). The subspecies *L*.*c. reissii* was finally represented by one paper only, so that it was not considered in the analyses.

### Statistical analysis

In order to perform parametric analyses, we first checked both the normality of the considered variables and the homogeneity of variances through the Levene’s test, transforming the variables that were not normal (Zar, 2009). Alternatively, we checked whether the kurtosis of these variables was positive, which allows us to assume a low probability of committing type I statistical error (Underwood, 1996).

We characterized the different types of culpeo diet by grouping the relative frequencies (RF) of the ten trophic groups into orthogonal factors carrying out a factor analysis and using the principal component analysis (PCA) algorithm. We also tested for spatial autocorrelation in the values of the extracted factors and the trophic diversity values (H’) by calculating Moran’s index I and their respective correlograms (Dormann et al., 2007; Rangel et al., 2010).

Pearson’s correlation between orthogonal factors and latitude, altitude, human footprint index and trophic diversity was calculated. In addition, we checked whether the latter correlated with each of the RFs of the trophic groups. We also explored whether the degree of protection of the study areas (with or without legal protection) and habitat type (scrub, steppe or mosaic) explained variation in the trophic factors. The influence of the presence of plant matter in the diet on trophic factors and the trophic diversity was also tested by performing a two-way and a one-way MANOVA analyses, respectively. We also assessed whether the presence of exotic lagomorphs, introduced into the study areas, influenced the diet of the species through a MANOVA with the trophic factors (excluding the factor that included the lagomorph group) plus the RF of big rodents and eggs. Furthermore, we analyzed the differences between the diets of the subspecies *L*.*c. andinus* and *L*.*c. culpaeus* by means of a MANOVA with the subspecies as a fixed factor and the trophic factors as dependent variables, as well as an ANOVA with the trophic diversity also as a dependent variable.

Finally, general linear models (GLM) were constructed for trophic diversity (H’) as a response variable, using 10 predictor variables: the four trophic orthogonal factors, latitude, altitude, protected area, habitat type, precipitation and the human footprint index. Of the total number of models obtained, the most parsimonious were identified through a selection process applying the Akaike’s criterion (Burnham & Anderson, 2002). Software for conducting the statistical analyses included SAM v.4.0 (Rangel et al., 2010) and Statistica 10 (StatSoft, 2011).

## Results

Relative frequency (RF) values of all the trophic groups calculated for each independent sample, as well as the Shannon-Wienner H’ trophic diversity index, are shown in Figure 2. The mean value of the latter was 1.23 (range: 0.36 - 1.88). The mean values of relative frequency (RF), expressed as a percentage, were: 41% small mammals, 21% lagomorphs, 12.4% invertebrates, 7.3% large herbivores, 7.3% birds, 6% big rodents, 2.9% herptiles, 0.9% carnivorous, 0.8% edentates and 0.7% eggs.

**Figure 2.**
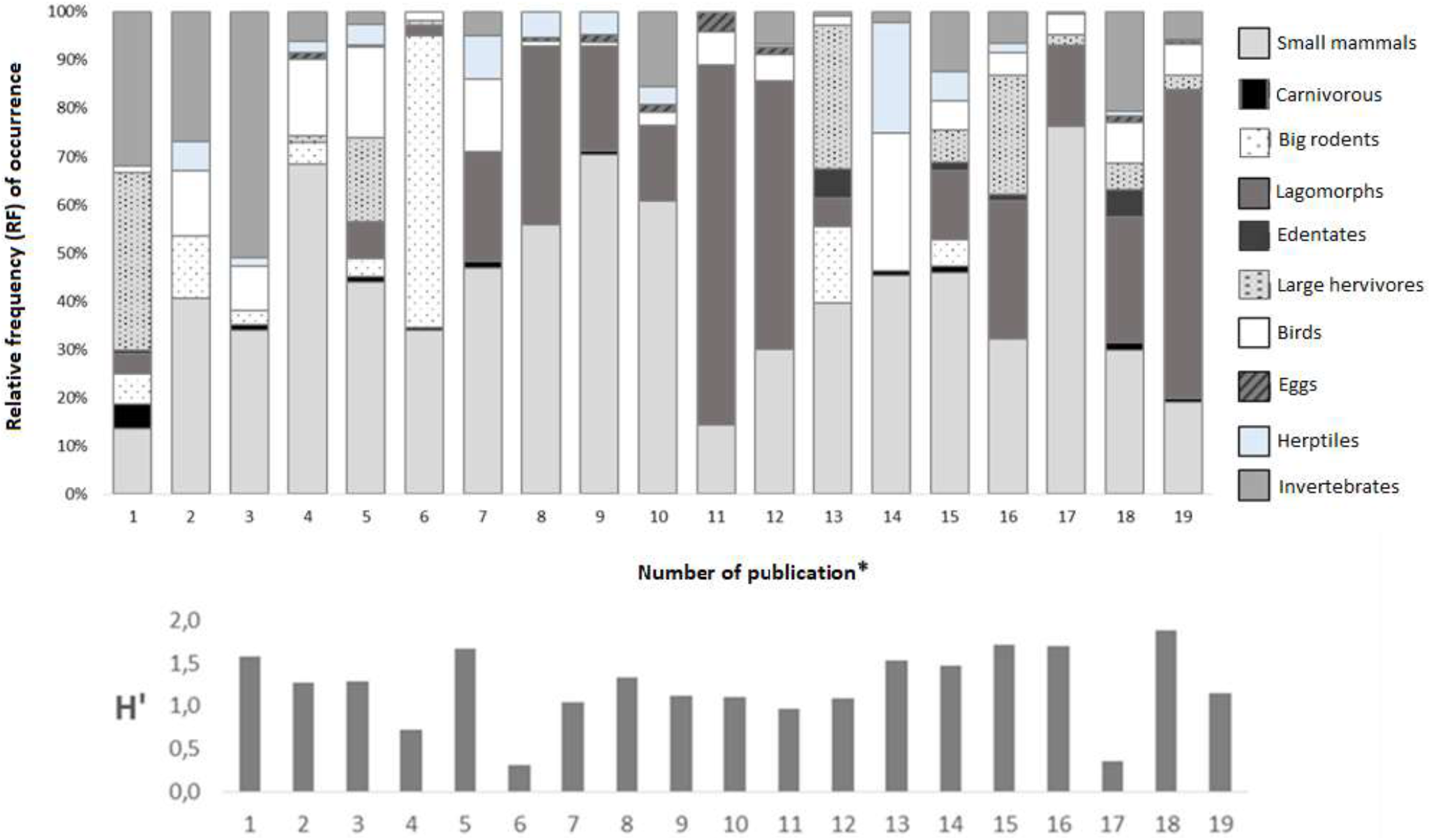
Relative frequency of occurrence (RF) of the ten trophic groups and values of Shannon-Wienner trophic diversity (H’) index for each independent sample considered in this study. * Publication number: 1. Guntiñas et al., 2017; 2. Romo et al., 1995; 3. Cornejo-Farfán & Jiménez-Milón, 2001; 4. Marquet et al., 1993; 5. Walker et al., 2007; 6. Pía et al., 2003; 7. Ebensperger et al., 1991; 8.9. Iriarte et al., 1989; 10. Jaksic et al., 1980; 11.12. Rubio et al., 2013; 13. Berg, 2009; 14. Achilles, 2007; 15. Palacios et al., 2012; 16. Novaro et al.,2000; 17. Monteverde et al., 2011; 18. Zapata et al., 2005; 19. Johnson & Franklin, 1994.

The factor analysis using the RF of the ten trophic groups generated four orthogonal factors (using standardized varimax rotation of axes) that overall explained 73.6% of the variance of the original variables (Table 1). The first factor describes a gradient from culpeo diets with high consumption of carnivorous, large herbivores and edentates (negative scores) towards less rich diets in these groups. The second factor is a gradient from high consumption of lagomorphs and eggs (negative scores) to poorer diets in these trophic groups and richer in big rodents (positive scores). The third factor is a gradient from diets with high consumption of birds and herptiles (negative scores) to low intake of these prey (positive scores). And finally, the fourth factor describes a gradient from diets with high consumption of small mammals (negative scores) to high consumption of invertebrates (positive scores).

**Table 1.** Description of the four orthogonal factors obtained from the factor analysis with the ten trophic groups considered. * indicates significant correlations of the variables.

| Variables | Factor<br>1 | Factor<br>2 | Factor<br>3 | Factor<br>4 |
| --- | --- | --- | --- | --- |
| Large herbivores | **-0.92\*** | 0.16 | 0.15 | -0.01 |
| Big rodents | 0.17 | **0.65\*** | 0.60 | -0.04 |
| Birds | 0.12 | 0.08 | **- 0.70\*** | -0.03 |
| Carnivorous | **-0.86\*** | 0.17 | 0.05 | 0.01 |
| Edentates | **-0.46\*** | -0.08 | 0.10 | 0.16 |
| Eggs | 0.21 | **-0.81\*** | 0.12 | -0.20 |
| Invertebrates | 0.14 | 0.32 | -0.43 | **0.74\*** |
| Herptiles | 0.27 | 0.25 | **-0.76\*** | 0.05 |
| Lagomorphs | 0.1 | **-0.9\*** | 0.33 | 0.13 |
| Small mammals | 0.30 | 0.13 | -0.21 | **-0.85\*** |
| Eigenvalue | 2.08 | 2.10 | 1.82 | 1.36 |
| **% Explained variance** | **20.82** | **21.00** | **18.18** | **13.57** |

According to the correlograms of the four trophic orthogonal factors and trophic diversity H’, which were based on Moran’s I index, the diet of the culpeo in South America was not spatially structured, since in general the values that describe its diet in the reviewed studies did not present spatial autocorrelation.

Factor 1 was negatively correlated with H’ trophic diversity (see Table 2), so the inclusion of carnivorous species, large herbivores and edentates in the diet of the culpeo significantly increased the trophic diversity of the canid, as well as the consumption of birds and herptiles, as factor 3 was also negatively correlated with trophic diversity. Factor 2 correlated positively with altitude, and negatively with the human footprint index and latitude (Figure 3). Therefore, at higher altitudes (and in areas of lower latitudes and human influence) the culpeo consumes more big rodents, while at higher latitudes and when the human footprint is greater in the environment, consumption of lagomorphs and eggs increases. Factor 4 was negatively correlated with altitude (Figure 3), indicating that higher altitudes result in higher consumption of small mammals and that invertebrates are more prevalent in the diet at lower altitudes. Taking each of the ten trophic groups one by one, those that contribute the most to the increase in the diversity of the culpeo diet are large herbivores and birds, while diets including big rodents tend to be less diverse (Table 2).

**Table 2.** Pearson’s correlations of the four trophic orthogonal factors extracted from the factor analysis with latitude, altitude, human footprint index and trophic diversity H’ index. The correlations between the latter and the relative frequency (RF) of the ten trophic groups considered in the review (n = 19) are also shown.

|  | Latitude | Altitude | Human footprint index | H' |
| --- | --- | --- | --- | --- |
| Factor 1 | 0.26 | -0.08 | 0.22 | -0.58 * |
| Factor 2 | -0.40 † | 0.58 * | -0.46 * | 0.13 |
| Factor 3 | 0.21 | -0.22 | 0.19 | -0.43 * |
| Factor 4 | 0.29 | -0.43 * | 0.02 | 0.25 |

|  | H' |
| --- | --- |
| Small mammals | -0.32 |
| Carnivorous | 0.33 |
| Big rodents | -0.42 † |
| Lagomorphs | -0.02 |
| Edentates | 0.54 |
| Large herbivores | 0.45 * |
| Birds | 0.17 * |
| Eggs | -0.17 |
| Herptiles | 0.29 |
| Invertebrates | 0.16 |

\* Significant correlation (p < 0.05). † Marginally non-significant correlation (p < 0.10)

**Figure 3.**
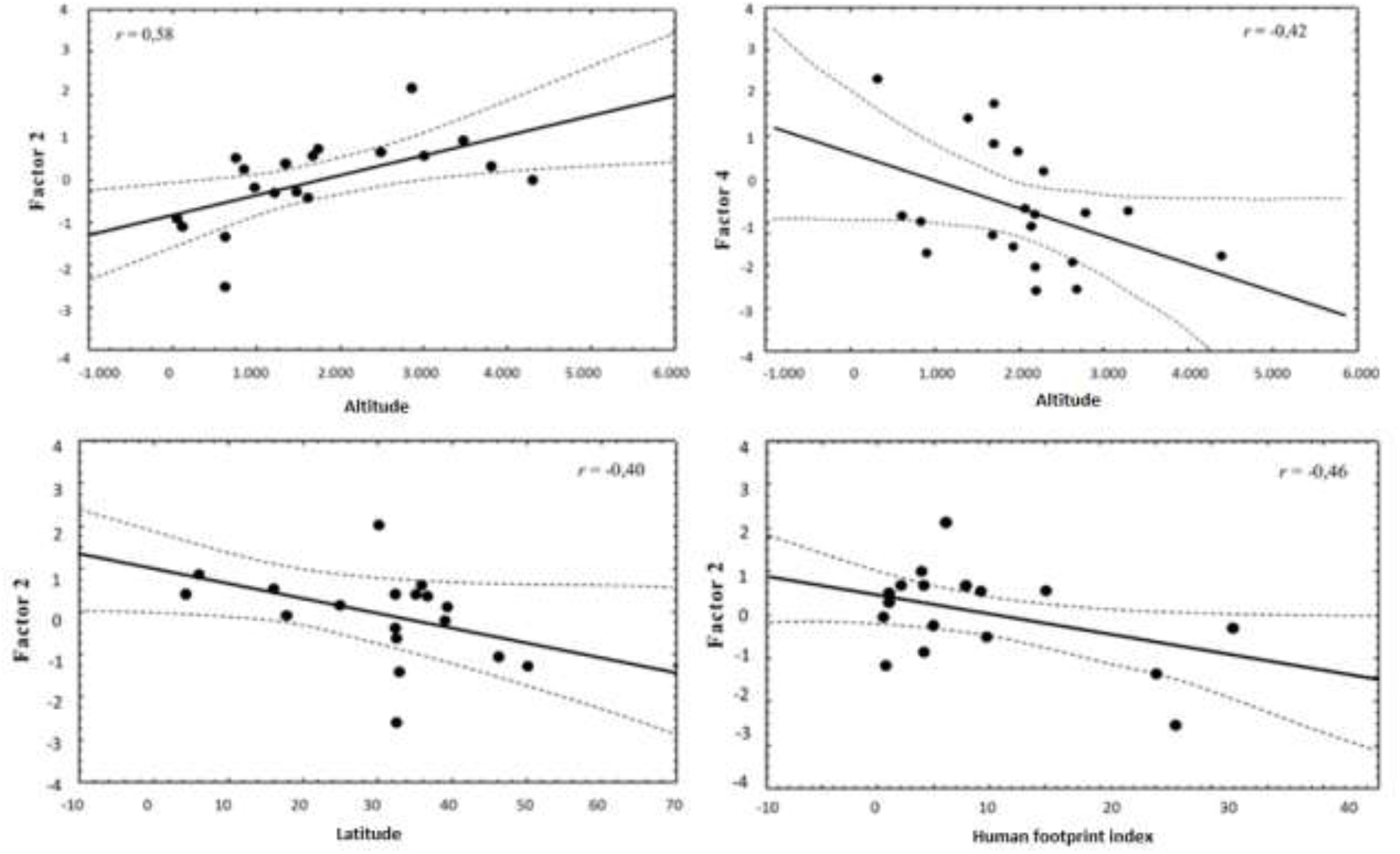
Relationships between factors 2 and 4 with altitude, as well as factor 2 with latitude and the human footprint index.

The consumption of plant material tended to influence the diet of the culpeo (MANOVA, Wilks’ Lambda = 0.59; F_4.14_ = 2.4; p = 0.09), with significant differences in factor 3 (F_1.17_ = 4.8; p = 0.04). Thus, when there was consumption of plants, there was also a greater consumption of birds and herptiles. However, the consumption of plant matter had no effect on the trophic diversity H’ index (F_1.17_ = 0.01; p = 0.98).

Habitat type influenced the diet of the culpeo given that significant differences were found in the trophic orthogonal factors (Table 3), specifically a positive relationship between factor 3 and the mosaic type habitat (F_2.13_ = 8.13; p = 0.005). Therefore, in mosaic habitats the culpeo consumed less birds and herptiles (Figure 4). No significant differences were detected in the trophic factors depending on whether the studies were carried out in protected or unprotected areas (Table 3).

**Table 3.** Two-way MANOVA with the four trophic orthogonal factors as dependent variables and two fixed factors (protected area and habitat type).

|  | Wilks'<br>Lambda | F | df<br>effect | df<br>error | p |
| --- | --- | --- | --- | --- | --- |
| Protected area | 0.63 | 1.49 | 4 | 10 | 0.28 |
| Habitat type | 0.25 | 2.53 | 8 | 20 | 0.04 |
| Interaction | 0.5 | 1.05 | 8 | 20 | 0.43 |

**Figure 4.**
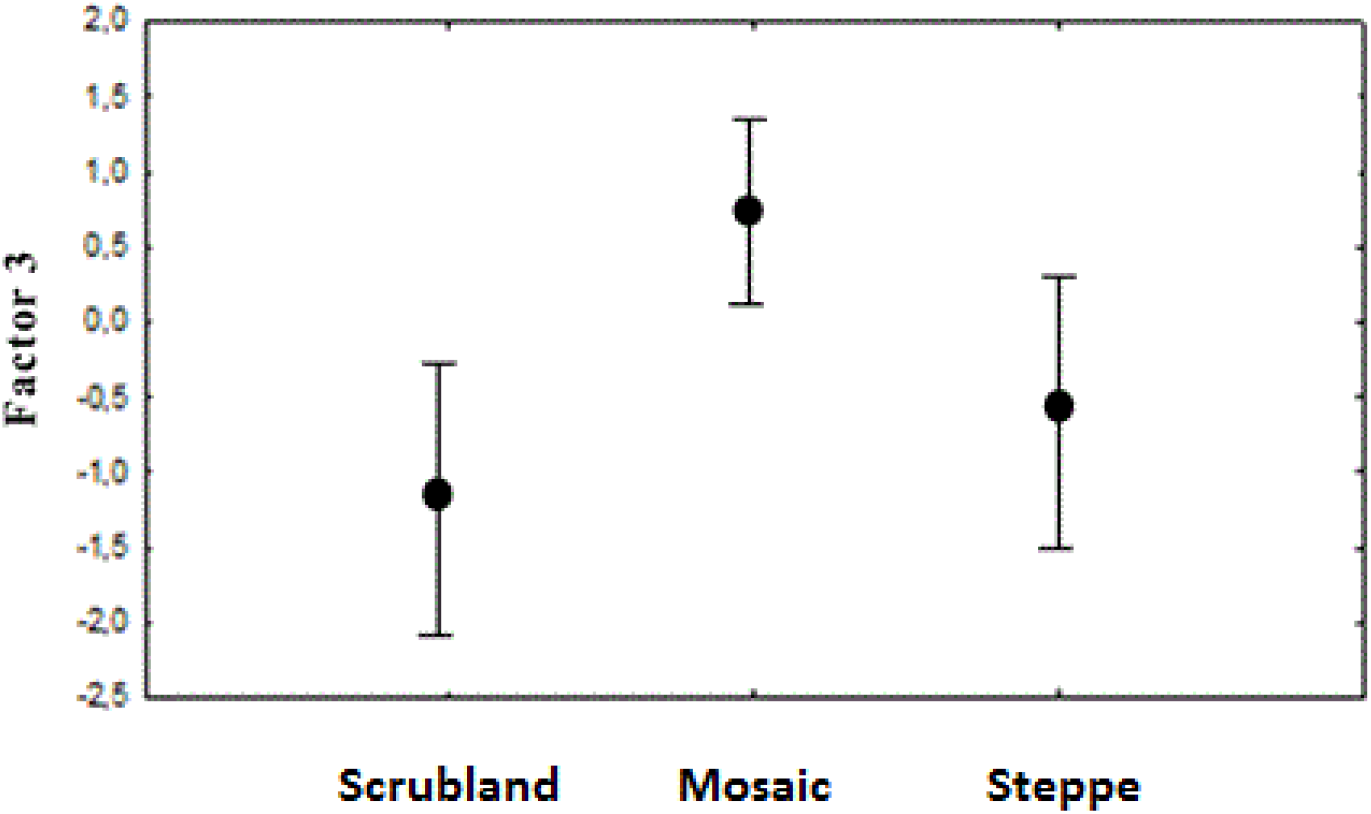
Relationship between factor 3 and habitat type: scrubland, mosaic and steppe (accompanied by standard deviation).

The presence of non-native (i.e. exotic) lagomorphs in the environment also influenced the diet of the canid (MANOVA, Wilks’ Lambda = 0.37; F_5.13_ = 4.46; p = 0.014), with a significant relationship with factor 3 (F_1.17_ = 10.41; p = 0.005). Thus, in areas with the presence of introduced lagomorphs the culpeos consumed fewer birds and herptiles (Figure 5).

**Figure 5.**
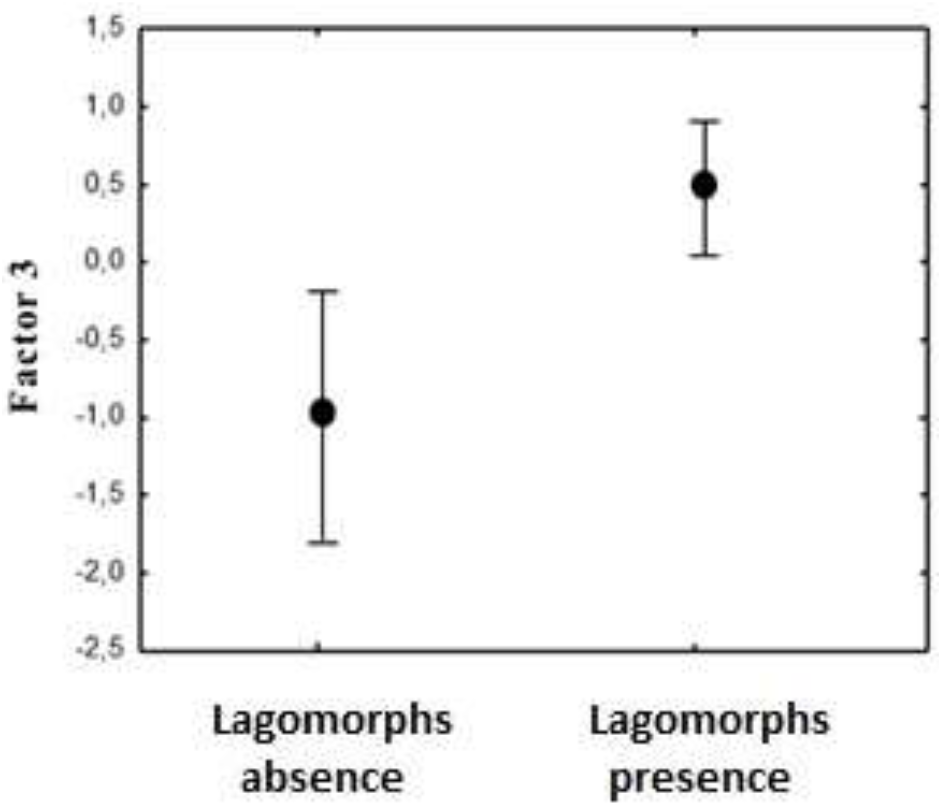
Relationship between the presence of exotic lagomorphs and factor 3 (accompanied by standard deviation).

Differences between subspecies *L*.*c. andinus* and *L*.*c. culpaeus* were not found in their diet, either considering the trophic orthogonal factors (MANOVA, Wilks’ Lambda = 0.37; F_4.13_ = 1.52; p = 0.25), or their respective H’ trophic diversity index (ANOVA, F_1.17_ = 0.03; p = 0.85).

In total, 1023 GLM models were generated for the trophic diversity (H’) as a response variable using the 10 predictors mentioned above. By applying the Akaike’s selection criterion only two models were found to be likely (Table 4): the most parsimonious (r^2^ = 0.75) incorporated factors 1 and 3 in addition to precipitation, while the next model (r^2^ = 0.79) also incorporated the protected area. In both models, the factors and rainfall were negatively correlated with the H’ trophic diversity index (Table 5), so that the most diverse culpeo diets were associated with areas of high rainfall and the consumption of carnivorous species, large herbivores, edentates, birds and herptiles (Figure 6). Considering the second most parsimonious model, culpeos that live in protected areas also show a greater diversity in their diet than those living in non-protected areas.

**Table 4.**
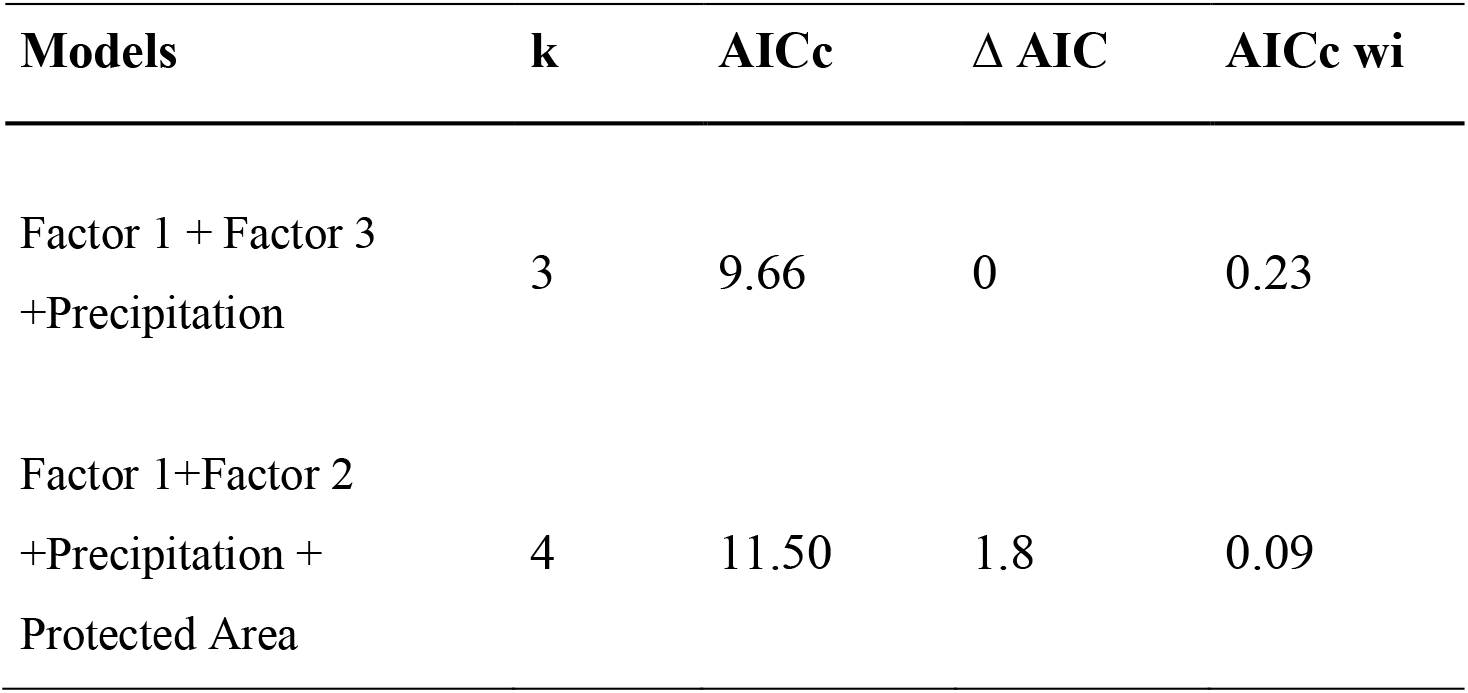
Explanatory models of culpeo’s trophic diversity (H’) index. It is shown the two parsimonious models obtained with the number of variables used (k), AIC values for small sample sizes (AICc), the difference between each selected model and the best model (Δ AIC) and each model weight (AICc wi). According to Burnham & Anderson (2002), only models with ΔAIC < 2 are really likely. The models were ordered from the lowest value (best model) to the highest value according to the AICc.

| Models | k | AICc | $\Delta$ AIC | AICc wi |
| --- | --- | --- | --- | --- |
| Factor 1 + Factor 3<br>+Precipitation | 3 | 9.66 | 0 | 0.23 |
| Factor 1+Factor 2<br>+Precipitation +<br>Protected Area | 4 | 11.50 | 1.8 | 0.09 |

**Table 5.**
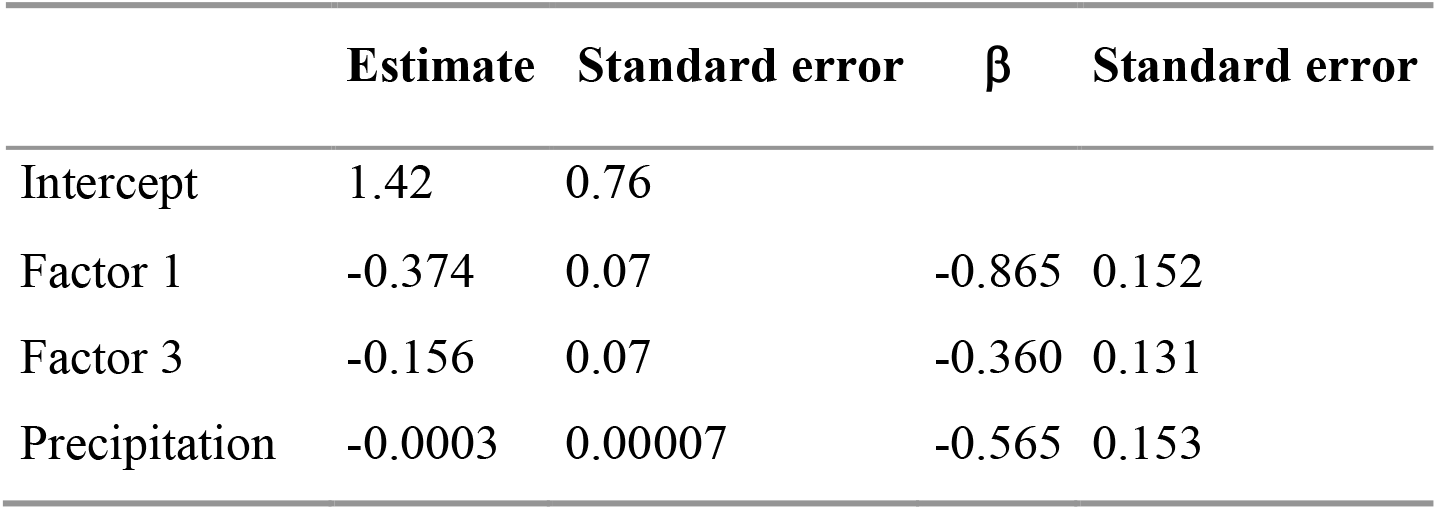
Estimates of the most parsimonious GLM model obtained to explain the variation in the trophic diversity (H’) index of the culpeo with factor 1, factor 3 and precipitation as predictors.

| | Estimate | Standard error | $\beta$ | Standard error |
| --- | --- | --- | --- | --- |
| Intercept | 1.42 | 0.76 |  |  |
| Factor 1 | -0.374 | 0.07 | -0.865 | 0.152 |
| Factor 3 | -0.156 | 0.07 | -0.360 | 0.131 |
| Precipitation | -0.0003 | 0.00007 | -0.565 | 0.153 |

**Figure 6.**
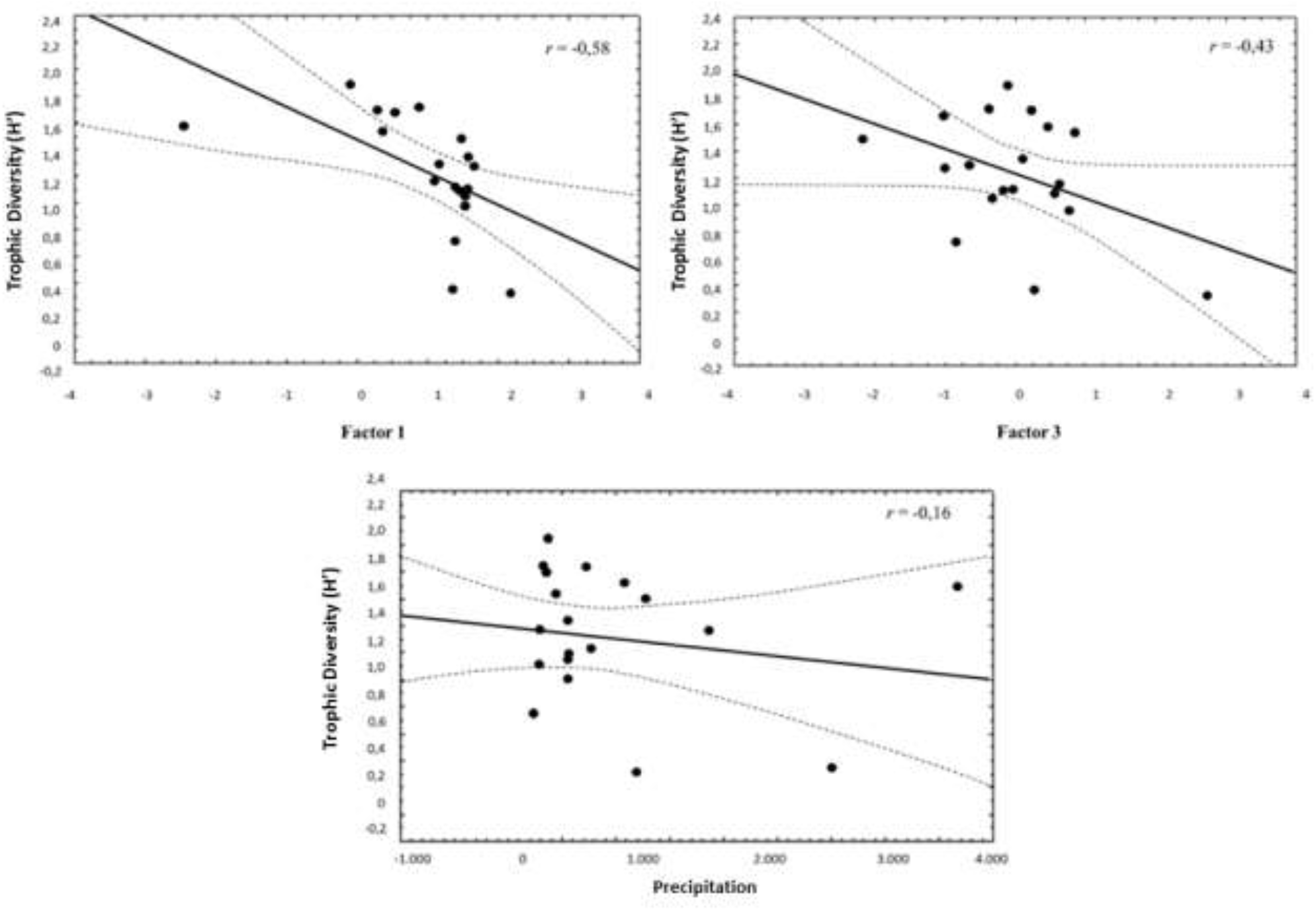
Relationship between trophic diversity (H’) index and factors 1 and 3, and precipitation, according to the most parsimonious GLM model obtained.

## Discussion

The culpeo is a medium-sized, mainly carnivorous canid (Redford & Eisenberg, 1992; Novaro, 1997a; Guntiñas et al., 2017), which can use a wide diversity of prey throughout its distribution range (Guntiñas et al., 2021). It has been described as a polyphagous species (Pía, 2011), as in some regions it uses trophic resources such as fruits and invertebrates (Ebensperger et al., 1991; Cornejo-Farfán & Jiménez-Milón, 2001; Guzmán-Sandoval, 2007). Thus, due to this varied diet between and within trophic groups, the culpeo has generally been considered as an opportunistic species (Crespo & De Carlo, 1963; Jacksic et al., 1980; Johnson & Franklin, 1994; Castro et al., 1994). To date, no dietary patterns have been described based on geographical variation, except for the exotic lagomorph consumption in the new areas the culpeo has expanded into, where native prey species seem to have been replaced by exotic ones (Lucherini, 2016; Guntiñas et al., 2021).

Our results show that small mammals are an important prey item across the culpeo’s range, as reflected in numerous studies (Iriarte et al., 1989; Ebensperger et al., 1991; Johnson & Franklin, 1994; Meserve et al., 1996; Novaro et al., 2000a; Pía et al., 2003; Zapata et al., 2005; Walker et al., 2007). In studies that quantified prey availability in the environment, a clear dietary selectivity has been repeatedly shown, i.e. in the face of a decline in small mammals (especially rodents), the diet did not change as might be expected in a truly opportunistic species (Jaksic et al., 1992; Martínez et al., 1993). In most of the studies reviewed small mammals have a high relative frequency of occurrence, although presenting a greater consumption at high altitudes. Therefore, the culpeo shows a certain degree of specialization, and could have a strong effect on small mammal populations and be a key predator regulating and/or limiting their populations (Krebs, 2002; Guntiñas et al., 2021).

Species are located at different levels within the food web, where top predators would be at the highest positions, potentially consuming the rest of the species, and with few or no other species that predate them (Essington & Hansson 2004; Essington et al., 2005). In this context, the pattern of diets rich in carnivorous species, large herbivores and edentates obtained in this review (associated with factor 1), clearly corresponds to those of an apical predator. In Andean systems, the only natural species that can prey on culpeos is the puma (*Puma concolor*) (see Guntiñas et al., 2021). Aside from the puma, the culpeo, also by preying on other carnivores, would act as an important regulator of mammal populations of species of equal or greater size (such as ungulates). This corresponds more to the role of a large carnivore than to those of small or medium-sized carnivores that the culpeo is classified into (Carbone et al., 2007), and indicates the key role of culpeos in high-Andean ecosystems (Guntiñas et al., 2021).

Regarding whether the presence of exotic lagomorphs introduced into the environment cause a change in the diet of the culpeo, the results of this review at the global scale indicate that where lagomorphs are present, there is a lower consumption of birds and herptiles. This pattern could be due to the reduced availability of these groups in environments where hares and rabbits are present, or because birds and herptiles are ignored in such areas (actually, these taxa appear to be secondary for the culpeo throughout its range). But beyond birds and herptiles, no widespread substitution of native prey (especially small rodents) has been detected by the dietary dominance of lagomorphs, as suggested in studies at the local scale (Crespo & De Carlo, 1963; Jaksic, 1998; Novaro et al., 2000a; Rubio et al., 2013), although in other studies this pattern was not observed (Meserve et al., 1987).

Considering the data globally, in the regions where lagomorphs are consumed, small mammals and other prey groups are still consumed, which were also important in terms of relative frequency of occurrence. It should be noted that the lagomorph prey group is the second most important trophic group in terms of RF intake. As the majority of these are exotic species (European hare *Lepus europeaus*, as well as the European rabbit *Oryctolagus cuniculus*), with only one study representing a native lagomorph, it is likely that the culpeo has taken advantage of this new environmental resource there where it has appeared, similarly to how it takes advantage of the availability of big rodents elsewhere. Furthermore, diets rich in lagomorphs and eggs as well as poorer in big rodents (according to factor 2), occur at low altitudes, low latitudes and in regions with an important human influence on the territory. Therefore, the consumption of lagomorphs, especially European hares (taking into account that they are also found at the highest latitudes), can be associated with lowlands and impacted populations of culpeos, such as agricultural and livestock areas, which may coincide with the works describing some substitution in the diet of native prey by exotic lagomorphs (Crespo & De Carlo, 1963; Jaksic, 1998; Novaro et al., 2000a; Rubio et al., 2013).

Depending on the altitude there is a clear pattern of trophic groups consumed, so that in higher altitude areas there is a greater consumption of small mammals and big rodents, while in lower lands there is a greater consumption of lagomorphs, eggs and invertebrates (Figure 7). In regions of low altitude, the high consumption of invertebrates appears to be consistent with some authors’ views on the importance of this group in highly seasonal Mediterranean ecosystems (Ebensperger et al., 1991; Correa & Roa, 2005). On the other hand, at high altitudes, the higher consumption of small mammals and big rodents could perhaps be due to their greater availability, a particular prey selection by culpeos, or also because of the lower availability of other trophic resources.

**Figure 7.**
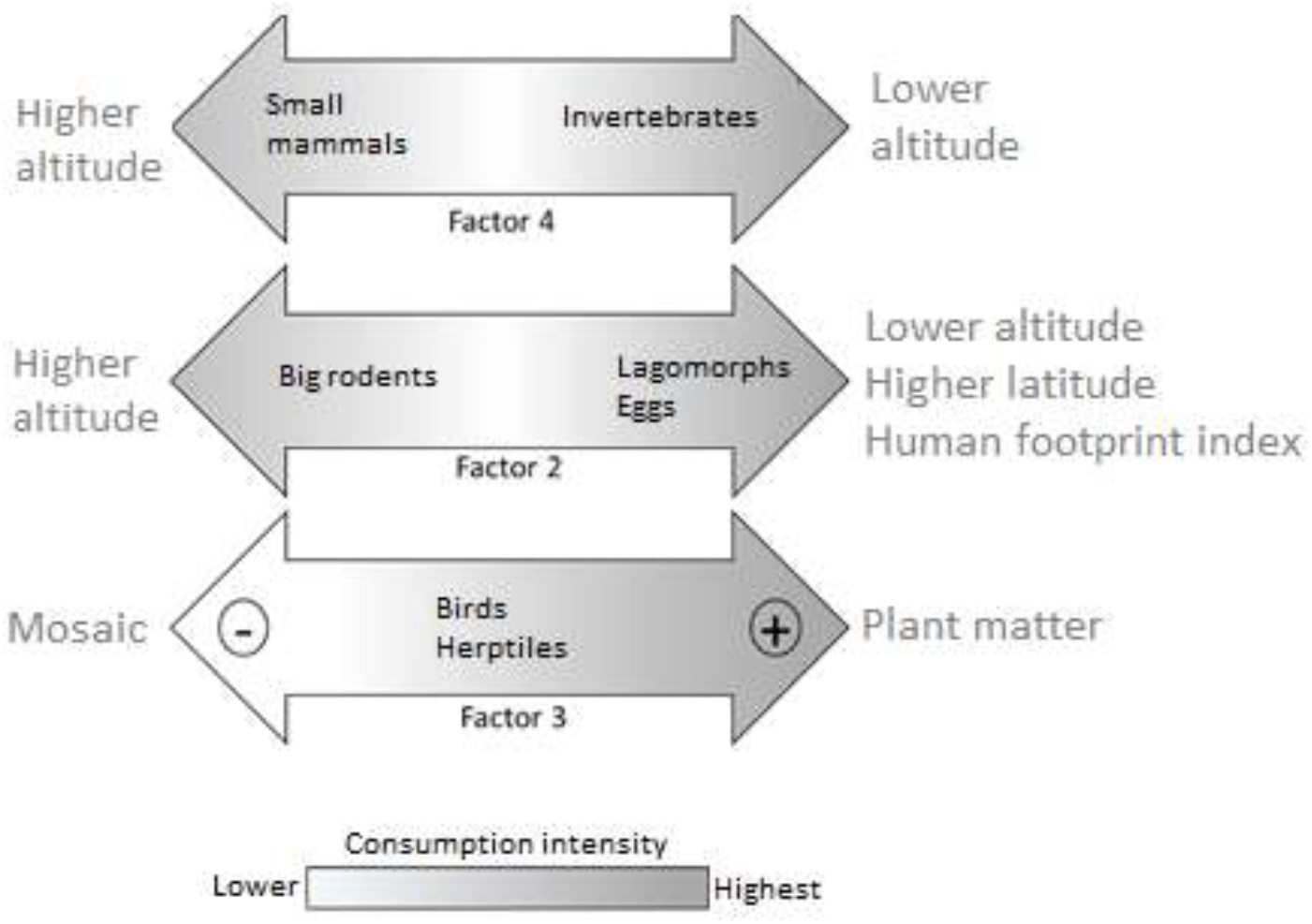
Conceptual model that describes the variation in consumption of trophic groups by the culpeo, grouped in orthogonal factors, in relation to some environmental variables according to the results obtained in this review.

It seems that the culpeo makes use of the resources available in each study area, determined by a series of environmental factors. Culpeos can specialize in them and so not behave merely as a generalist or opportunist. The aforementioned evidence that culpeos can maintain their consumption of small mammals even when their abundance decreases, or the high degree of consumption of large herbivores in certain regions as is the case with ungulates in the high Andes of Ecuador (Guntiñas et al., 2017), indicates that the culpeo behaves more like a facultative specialist than a strict generalist (Glasser, 1984; Guntiñas et al., 2017).

When culpeos consume plant material they often include other alternative or secondary food resources, such as birds and herptiles (Figure 7), perhaps also in response to the low abundance or availability of preferred prey (such as rodents) and therefore due to the need to supplement the diet. In addition, the consumption of birds and herptiles was lower in mosaic-type habitats, indicating a possible reduced availability for the culpeo of this prey in scrubland and steppe areas.

No differences have been found in the diet between the subspecies *L*.*c. andinus* and *L*.*c. culpaeus*, neither in the trophic groups consumed nor in the trophic diversity, suggesting that both subspecies display a similar trophic behavior. However, as more diet publications emerge in regions where the ecology of culpeo is poorly known, this picture could change. So, it would be interesting to review the trophic data again in the future, including also the subspecies *L*.*c. reissii* and increasing the number of case studies of *L*.*c. andinus*.

On the other hand, one of the best-known biogeographical patterns of dietary variation in a number of species is that it is a function of the latitudinal gradient, increasing from the poles towards the tropics (Rosenzweig, 1995). Altitude is also an important driver of species diversity in the diet, as mountains are sites of high diversity and endemism, although under particular conditions this pattern is not always met (Gentry, 1995). In the case of generalist predators with a wide range of distribution, whose diets reflect the availability of prey in the environment, changes would be expected according to these biogeographical patterns (e.g. Clavero et al., 2003; Lozano et al., 2006). In the case of the culpeo, with populations distributed in a marked latitudinal (from 3° to 50° south latitude) and altitudinal (from sea level to 3,800 m) gradient, no differences in trophic diversity were found based on these geographical variables. This result could be explained because the culpeo does not behave like a generalist predator, as mentioned above, so its role as a facultative specialist carnivore (Guntiñas et al., 2017, 2021) could mask latitudinal and altitudinal effects by specializing locally in particular trophic resources. According to the most parsimonious model obtained, the most diverse culpeo diets are those associated with high precipitation rates, as well as the consumption of carnivorous species, large herbivores, edentates (i.e. armadillos), birds and herptiles (Figure 8). In addition, it is also possible that in areas with some type of government protection the canid’s diet may be more diverse. Well-conserved areas, which host populations of species such as large herbivores, carnivores and edentates, would presumably have a greater diversity of species, and therefore of potential prey types, that culpeos could incorporate into their diet.

**Figure 8.**
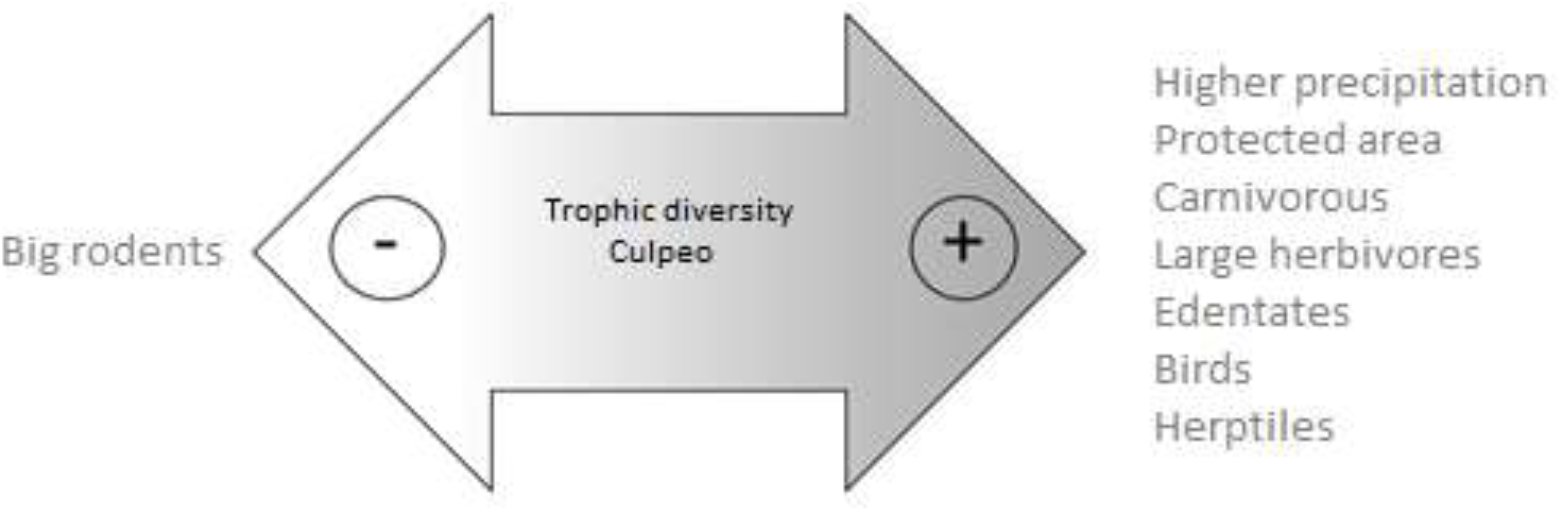
Conceptual model describing the variation in trophic diversity (H’) of the culpeo diet according to the results obtained in this review.

## Acknowledgments

This research was carried out partly with the economic support of the Universidad Técnica Particular de Loja (UTPL), Ecuador. JL was supported by a Prometeo Fellowship from SENESCYT, the National Agency for Education and Science of Ecuador, between 2014 and 2015. He was also supported by Department of Biodiversity, Ecology and Evolution in Complutense University of Madrid (Spain) during the editing of this article. AFM was supported by a Ramón y Cajal research contract from the MINECO (RYC-2016-21114).

